# GPU accelerated partial order multiple sequence alignment for long reads self-correction

**DOI:** 10.1101/2020.02.14.946939

**Authors:** Francesco Peverelli, Lorenzo Di Tucci, Marco D. Santambrogio, Nan Ding, Steven Hofmeyr, Aydın Buluç, Leonid Oliker, Katherine Yelick

## Abstract

As third generation sequencing technologies become more reliable and widely used to solve several genome-related problems, self-correction of long reads is becoming the preferred method to reduce the error rate of Pacific Biosciences and Oxford Nanopore long reads, that is now around 10-12%. Several of these self-correction methods rely on some form of Multiple Sequence Alignment (MSA) to obtain a consensus sequence for the original reads. In particular, error-correction tools such as RACON and CONSENT use Partial Order (PO) graph alignment to accomplish this task. PO graph alignment, which is computationally more expensive than optimal global pairwise alignment between two sequences, needs to be performed several times for each read during the error correction process. GPUs have proven very effective in accelerating several compute-intensive tasks in different scientific fields. We harnessed the power of these architectures to accelerate the error correction process of existing self-correction tools, to improve the efficiency of this step of genome analysis.

In this paper, we introduce a GPU-accelerated version of the PO alignment presented in the POA v2 software library, implemented on an NVIDIA Tesla V100 GPU. We obtain up to 6.5x speedup compared to 64 CPU threads run on two 2.3 GHz 16-core Intel Xeon Processors E5-2698 v3. In our implementation we focused on the alignment of smaller sequences, as the CONSENT segmentation strategy based on k-mer chaining provides an optimal opportunity to exploit the parallel-processing power of GPUs. To demonstrate this, we have integrated our kernel in the CONSENT software. This accelerated version of CONSENT provides a speedup for the whole error correction step that ranges from 1.95x to 8.5x depending on the input reads.

## I. Introduction

Third generation sequencing technologies, such as Pacific Biosciences (PacBio) and Oxford Nanopore Technologies (ONT), are establishing themselves as effective tools for important genomic problems such as assembly. They provide much longer reads compared to the second generation technologies such as Illumina, which allows a more precise contig and haplotype assembly and structural variant calling. However, third generation reads have much higher error rates compared to their second generation counterparts. Although the error rate of these sequences is improving, they are still in the 10-20% range, which is significantly higher than 0.2% error rates of Illumina sequencing. Moreover, third generation sequences present a more complex error profile with a high prevalence of insertions. These factors justify the need to include error correction as a preliminary step in any genome analysis project that uses third generation sequences. In recent years, many different tools and methodologies have been employed to tackle this problem [1]. Due to the wide availability of third generation sequences and the improvements achieved in reducing their overall error rate, self-correction methods are becoming the preferred approach as opposed to hybrid methods, which rely on second generation sequences to correct the noisy reads.

A commonality among many of these self-correction tools is that they rely on multiple sequence alignment (MSA) to obtain a consensus sequence to provide corrections for the raw reads. In particular, tools such as RACON [2] and CONSENT [3] use partial order (PO) alignment to generate consensus sequences. PO alignment can be viewed as an extension of the traditional Needleman-Wunsch (NW) and Smith-Waterman (SW) algorithms for optimal sequence alignment, that interprets sequences as partially ordered graphs and allows to encode the intermediate results of the MSA more completely and effectively. This comes with the additional computational cost of dealing with graphs instead of sequences and requires a more generalized algorithm to compute the globally optimal alignment result.

In this paper, we present a GPU implementation of a PO MSA algorithm, based on a fork of the POA V2 software library [4]. Lee et al. [5] described the general concepts behind this PO alignment library. We verify that our implementation optimally leverages the GPU by using an extended Roofline model analysis. In addition, we present an integration of our kernel with CONSENT, a state of the art self-correction tool. While our work is motivated by error correction, MSA in general and PO alignment in particular have widespread applications in computational biology that we summarize in Section II. The techniques we describe in this paper will help others develop optimal parallel GPU implementations of similar algorithms in the MSA family. The main contributions of this work are:

- A GPU implementation of the PO alignment algorithm that achieves up to 6.5x speedup compared to the software version run on two 2.3 GHz 16-core Intel Xeon Processors E5-2698 v3 with 64 CPU threads.
- An extension of the Roofline model analysis for GPUs presented in [6] that takes into account the characteristics of GPU architectures and algorithmic features of alignment algorithms to evaluate the performance of our implementation on the NVIDIA Tesla V100 GPU.
- The integration of our kernel with CONSENT, a state of the art long read self-correction tool, that employs a more optimal software infrastructure to fully utilize the computation capabilities of the GPU, obtaining up to 8.5x speedup of the error correction module without affecting the quality of the results. Both software are run on two Intel Xeon Gold 6148 (‘Skylake’) running at 2.40 GHz with 80 software threads and an NVIDIA Tesla V100 GPU.

The remainder of the paper is organized as follows. In Section II, we discuss similar works and how they relate to our implementation of PO alignment. In Section III, we report the key concepts behind the PO alignment algorithm for the reader’s convenience. In Section IV, we detail the GPU implementation of our alignment kernel, the strategies employed to obtain maximal acceleration and the reasoning behind them. In Section V, we explain how the kernel was integrated with CONSENT. In Section VI, we present the results obtained by testing our implementation. Finally, Section VII summarizes our contributions and details possible future research directions for this work.

## II. Related work

Multiple sequence alignment (MSA) is an extremely important task in computational biology. MSA is performed on a group of related protein, DNA or RNA sequences and aims to find the globally optimal alignment between these sequences that maximizes a given scoring function. For example, profile search methods such as PSI-BLAST [7] and profile hidden Markov models [8] that form the backbone of the most accurate protein homology search algorithms first start with the MSA of the set of proteins that are already known to be homologous to each other.

Since the general MSA problem is known to be NP-hard [9], several heuristic approaches have been developed to render the task more manageable. Progressive MSA reduces the problem of multiple sequence alignment to a series of iterated pairwise alignments. Clustal-W [10] adopts this approach paired with other heuristics to obtain improved sensitivity. Moreover, a GPU-accelerated version of this algorithm has been shown to achieve notable speedup [11]. Another example of GPU acceleration following the same approach is due to [12]. The T-coffee alignment algorithm [13] is another example of heuristic approach that has been adopted to improve the accuracy of results for pairwise multiple sequence alignment. This approach has also been accelerated on a GPU architecture [14].

While these approaches based on the pairwise alignment of sequences have useful applications, in the context of self-correction the encoding of the result of multiple sequence alignment as a single string is not optimal to obtain a final consensus sequence. For this application, PO alignment achieves notable results since it can model more complex alignment structures [15]. Variations of the PO alignment approach has also been the workhorse of many popular whole genome alignment tools [16], [17]

Since handling partial order graphs introduces an additional element of complexity to the traditional dynamic programming alignment algorithms, a GPU acceleration of this task is more challenging. To the best of our knowledge, the only other GPU implementation of PO alignment has been recently presented by NVIDIA as a part of the Clara Genomics Analysis SDK [18]. A more detailed comparison with our work is discussed in section VI.

### III. Background

The Partial Order Multiple Sequence Alignment (PO-MSA) algorithm extends the dynamic programming method of Needleman-Wunsch for the alignment of two sequences. In PO alignment the input sequences become partial orders containing branching. Each node in the partial order represents a base, which is connected to other bases by directed edges. These partial order structures are transferred to the two-dimensional dynamic programming matrix by flattening the nodes to a sequence that is a legal ordering according to the original partially ordered graph while maintaining the edges of the original structure. To calculate the score of each cell the NW algorithm admits only three possible moves, insertion, deletion or substitution. In PO alignment this move set is extended since each cell can have more than one predecessor both vertically, horizontally and diagonally in the DP matrix depending on the structure of the input PO graphs. The scoring system associates a transition cost to each move, depending on whether the move is diagonal, vertical or horizontal according to the selected match, mismatch, and gap penalties. For a cell *S* of the DP matrix aligning residues *n* and *m* with gap penalty *g*, let *w*(*n, m*) be the match or mismatch score for a diagonal move, *p*(*n*) the horizontal predecessor of a cell *S, q*(*m*) its vertical predecessor. The scoring is calculated as:

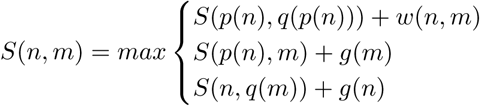

for each vertical and horizontal predecessor of the involved cells. The iterative MSA procedure starts by encoding the first two sequences as trivial partial orders. The PO alignment algorithm is then applied to obtain the optimal alignment between the sequences. The alignment result is used to fuse the two original sequences, as illustrated in figure 1. Where the sequences align, fuse the nodes of the two partial orders. Where they do not align, create a branch in the partial order and keep both fragments of the two sequences. We then perform PO alignment on the fused partial order graph and iteratively enrich it with the results of the alignment to the other sequences until we obtain a final PO graph that encodes the result of our MSA procedure.

**Fig. 1.**
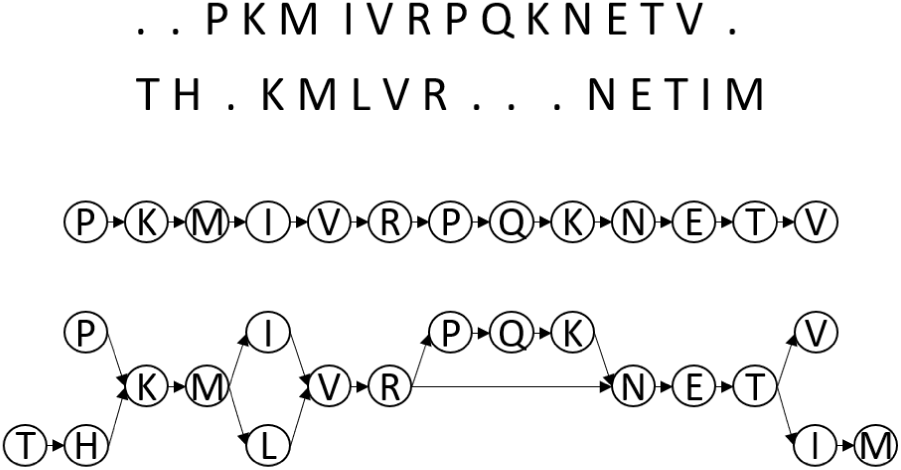
Interpretation of a sequence as a partial order graph and corresponding fused graph obtained from the alignment result of two sequences

## IV. Implementation

### A. Kernels Organization

Our GPU implementation of the PO-MSA of the BOA library is logically divided into a few specialized kernels that interact with each other to compute a batch of alignments. The input consists of a batch of windows of sequences to align and the final output is for each element in the batch the corresponding MSA. Each GPU block in each kernel takes care of processing one MSA task in the batch. At the beginning of execution, a portion of the GPU global memory is reserved depending on the dimension of the input sequences. This statically allocated memory space is empirically determined to be sufficient to complete the whole MSA procedure.

Nevertheless, the dimension of the MSA problem is data dependent, as the memory footprint of the partial order which encodes the alignment result depends on how the sequences in the window align. We have avoided considering the worst case space for the PO graph, as doing so would severely impact the kernel performance. Instead, since most alignments require a far smaller amount of space when the sequences in the window align well to each other, we allocate an empirically determined amount of space and in the rare examples where this space is not sufficient, the kernel is able to default to the CPU implementation of the algorithm to obtain the correct result. Since this instance is very rare, the overall impact on performance is negligible. Once the memory on the GPU is reserved, the *generate lpo* kernel is called. This kernel transforms the sequences in each window into linear partial order graphs. This operation is performed in parallel for each sequence in the window, and each window in the batch. After this initialization step is complete, the main alignment loop executes. For each pairwise alignment to perform, the following steps are executed:

- *Initialization*: the DP matrices are initialized for the next pair of alignments in the batch.
- *Pairwise PO alignment*: the actual alignment kernel is executed. This kernel performs the scoring and backtracking phases in parallel for each element in the batch.
- *PO graph fusion*: the result of the pairwise alignment is used to integrate the sequence aligned in the previous step to the current PO graph encoding the MSA result

Once all the sequences in the window have been processed for all the elements in the batch, the final alignment result is computed for each element and transferred back to the CPU.

### B. Scoring kernel Optimization

Since most of the PO graphs handled during the MSA procedure are very sparse, we chose to represent them as a sequence of characters which encode the bases contained in the nodes and an edge list containing the predecessors to each node represented by their position in the sequence. Since we want to avoid dealing with actual lists on the GPU, we use three arrays: the *sequence* array contains the characters of the PO graph nodes in an admissible total order consistent with the PO graph. The *edge_list* array contains all the edges for the graph stored in contiguous memory space and the *edge_offsets* array contains the end positions of each set of predecessors for each node in the PO graph. To efficiently access multiple PO graphs in parallel while processing a batch, we also store global offsets indexing the start of each array for each element in the batch which enables us to store all the PO graphs in contiguous memory.

One of the most challenging aspects of this generalized scoring algorithm is that we need to access a variable number of memory cells at runtime to compute the correct score for a cell. To reduce the global memory accesses caused by accessing the auxiliary *edge_list* and *edge_offsets* arrays we transfer these structures in the GPU shared memory before scoring. To parallelize the scoring process as much as possible, we compute the score for each cell in each antidiagonal in parallel. This is done by assigning a GPU thread to each column of the DP matrix. At each step of the scoring process a thread takes care of scoring a cell in its column starting from the top, and each thread is activated a step after the one to its left, as shown in figure 2. This exploitation of parallelism in the algorithm is still possible even if we are dealing with graphs instead of sequences, because the PO graphs are directed and acyclic, which ensures that a predecessor for any given cell in the DP matrix must be somewhere in the triangular up-left corner delimited by the antidiagonal which is currently being processed.

**Fig. 2.**
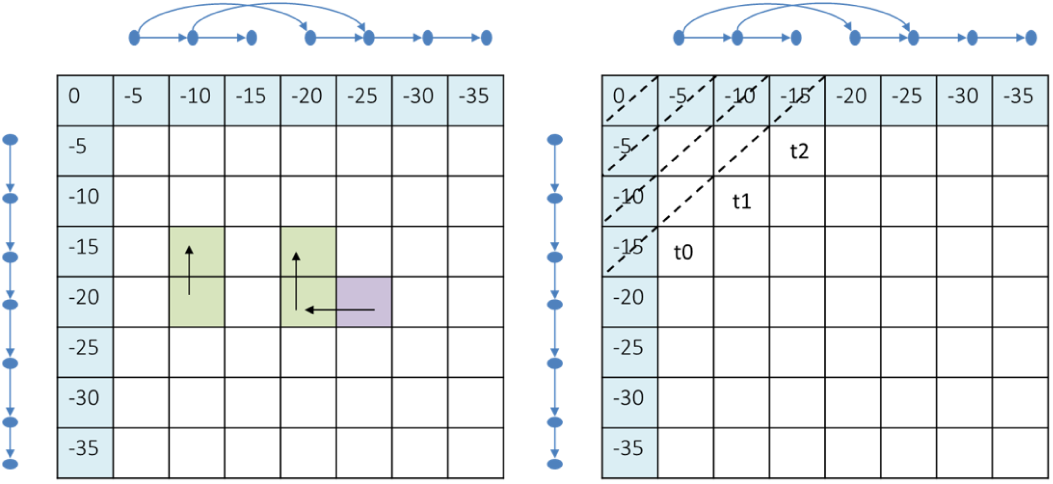
Computation of the dynamic programming matrix for a PO alignment. In the first image the data dependencies for a scoring cell are highlighted. The second image highlights the wavefront direction as well as an example of thread assignment for the GPU kernel

Nevertheless, this generalization creates an important problem: in the traditional SW and NW algorithms it is enough to store the two previous antidiagonals of the DP matrix and the current maximum score to complete the scoring of a new antidiagonal. In PO alignment, each antidiagonal has a set of dependencies which vary depending on the topology of the two PO graphs that we need to align, and lie anywhere in the top-left corner of the DP matrix. In the original software library, a bookkeeping mechanism is used to free columns of the DP matrix where all their cells are not reached by any node ahead in the computation. In our GPU implementation we tried to employ a similar mechanism, but ultimately removed it since it did not yield any performance improvement and for our target use case it was possible to store the whole DP matrix in the GPU global memory. Nevertheless, we are aware that for a different GPU with less memory resources, or to compute alignments for larger graphs, some heuristic procedure to reduce the memory footprint may be necessary, and it is a possible improvement to our approach.

To obtain coalesced memory access for each antidiagonal in the DP matrix, we store it in global memory one antidiagonal after another. This allows for an optimal memory access pattern for each thread and simplifies the indexing mechanism during the computation. To easily access the correct cells we need to store an array containing the offsets of the first cell of each antidiagonal. The dimensions of each pairwise alignment in the MSA procedure changes at each iteration and it is data dependent, therefore we need to compute the correct offsets at each iteration. The offsets can be easily deduced from the dimensions of the DP matrix and we can compute them without a significant impact on performance.

We use an additional pair of matrices, *move*_*x*_ and *move*_*y*_, to store the DP moves by which a cell of the DP matrix has been assigned its score. It is important to note that, differently from the traditional NW and SW algorithms, we cannot simply store the direction of the predecessor used for the scoring of a cell, but we need to store its coordinates: if cell *c* has been scored using the value of cell *c*2, *moves*_*x*_ [*c*] = *c.x* − *c2 .x*, and *moves*_*y*_ [*c*] = *c.y* − *c2 .y*, where .*x* and .*y* are the position of the cell in the DP matrix. The maximum score is calculated for each column in the DP matrix and stored by each thread in a shared array, together with the coordinates of the cell it originates from. At the end of the scoring procedure, the final maximum score is calculated via a parallel wrap reduction on the shared array. The backtracking procedure is executed by a single thread, but it contributes in minimal part to the overall execution time of the scoring procedure (around 1% of the overall execution).

### C. Roofline Model Analysis

In this section, we provide a detailed performance analysis of our PO alignment kernel by adapting the Instruction Roofline model [6]. The Instruction Roofline model is a visually-intuitive method to understand the performance of a given kernel based on a *bound and bottleneck* analysis approach. The Instruction Roofline model characterizes a kernel’s performance in billions of warp instructions per second (warp GIPS) as a function of its instruction intensity (II, x-axis, warp instructions per DRAM memory transaction) on GPUs. Given that our PO alignment kernel performs only integer instructions, here we combine billions of warp integer instructions (warp GIntIPS), instruction intensity (warp integer instruction per transaction), and memory bandwidth into a 2D log-log scale graph. A given implementation of the target application is represented as a point on the graph. If the point touches the horizontal ceiling, we know that we are compute bound. Conversely, if it touches the memory bandwidth (the oblique line), we know that the application is memory bound. If the point does not touch either line, it is possible that by using more resources, either bandwidth or computational power, the application could be further optimized.

To verify the optimality of our GPU implementation, we propose a tighter ceiling which is derived from a study of the parallelism characterizing a particular algorithm. In every alignment algorithm solved through dynamic programming we have two fundamental dimensions of parallelism, intertask and intra-task. Inter-task parallelism can be achieved on GPU by scheduling multiple alignments tasks at the same time, while intra-task parallelism is achieved by computing each antidiagonal of the DP matrix in parallel. This intra-task parallelism limits by nature the number of cells of the DP matrix that we can compute each cycle for a single alignment. Since the number of GPU threads scheduled per block is fixed for a given kernel execution, if we assign the computation of each cell of the DP matrix to a thread to achieve maximum parallelism, at the beginning of the computation we will have several inactive threads. This performance loss can be mitigated by reducing the number of parallel threads in favor of scheduling more blocks computing more alignments at the same time. In this context, we consider an implementation to be optimal if the frequency of integer operations reflects the maximum achievable frequency given the number of active elements expected at a given cycle of the algorithm and the number of threads and blocks scheduled. This indicates that a given implementation is not slowed by memory latency issues, which given the complex nature of the memory accesses in our algorithm, is a critical issue to evaluate. This ceiling is computed as:

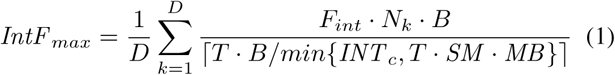

Given a set of alignment problems chosen to evaluate the kernel, *D* is the average number of antidiagonals of the DP matrices, *F*_*int*_ is the frequency of an integer functional unit on the GPU, *N*_*k*_ is the number of cells processed at each step for the average alignment and *B* is the number of scheduled GPU blocks. *T* is the total number of threads required to compute *N*_*k*_ cells, computed as 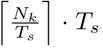, where *T*_*s*_ is the number of thread scheduled per block when the kernel is called. *INT*_*c*_ is the number of integer functional units available on the NVIDIA Tesla V100, *SM* is the number of streaming multiprocessors and *MB* is the maximum number of blocks per streaming multiprocessor.

This model captures the fact that the number of overall scoring cells computed at each iteration is limited by the maximum parallelism of the algorithm at the given iteration, the number of blocks and threads scheduled and the maximum physical resources of the GPU. We assume that an optimal implementation can fully utilize the integer functional units available to each thread which is actively computing the scoring for a DP matrix cell at each iteration. This ceiling is necessarily lower than the more theoretical ceiling which assumes that every thread is always able to compute a valid DP cell. Of course this result is achievable in theory by assigning only one thread to each alignment problem, but this is not achievable in practice since CUDA issues instructions at the warp level, which is constituted by 32 thread. Moreover, the optimal balance between inter-task and intra-task parallelism also depends on the processing overhead generated by the segmentation of the single tasks if we use fewer threads than the size of the maximum antidiagonal. This model is useful to verify that a chosen distribution of these two level of parallelism is using optimally the technological resources available.

In Figures 3, 4, 5 and 6 we have plotted the result of our Roofline analysis for four of the combinations of scheduled threads and blocks used in the kernels parameterized for windows of sequences with maximum initial length of 32, 64, 128 and 255 bp respectively. On the y axis we have the warpgiga-instructions per second, on the x axis we have the warp instructions per memory transaction and a single point on the graph represents an instruction intensity. The diagonal lines represent the different bandwidth for global memory, L1 and L2 caches, and the ceilings represent, from highest to lowest, the maximum theoretical ceiling, empirical ceiling for integer instructions and our proposed theoretical ceiling in terms of warp-giga-integer-instructions per second. We observe that in all four cases the memory bandwidth is not the limiting factor. As we increase the number of parallel threads scheduled, the value of the integer operational intensity, marked by the triangular dots in the graph, increases. This is primarily because the average size of the alignment problem increases as the length of the sequences to align increases, therefore the percentage of time effectively spent computing the scoring matrix increases with respect to the initial time spent in the setup phase and the final backtracking phase. Since we have taken the average execution time of the alignment kernel to compute the operational intensity, the overall operational intensity increases proportionately. In all the figures we have the integer operational intensity near our predicted theoretical ceiling, which is matched in figure 6. This tells us that the kernel is optimally using the integer functional units for the given number active threads and scheduled alignments.

**Fig. 3.**
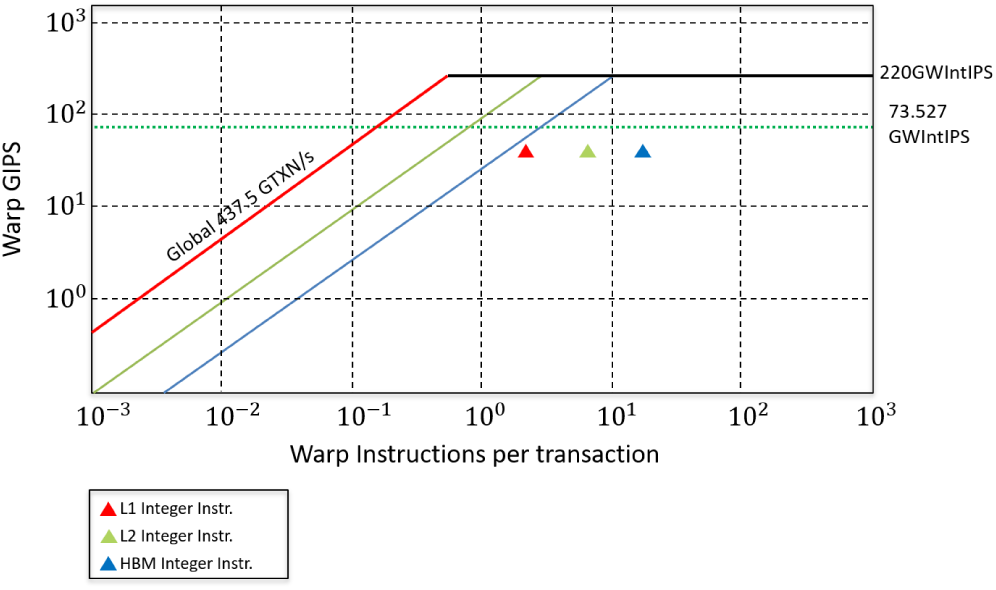
Roofline model plot for a kernel with 32 threads per block and 150000 parallel alignments

**Fig. 4.**
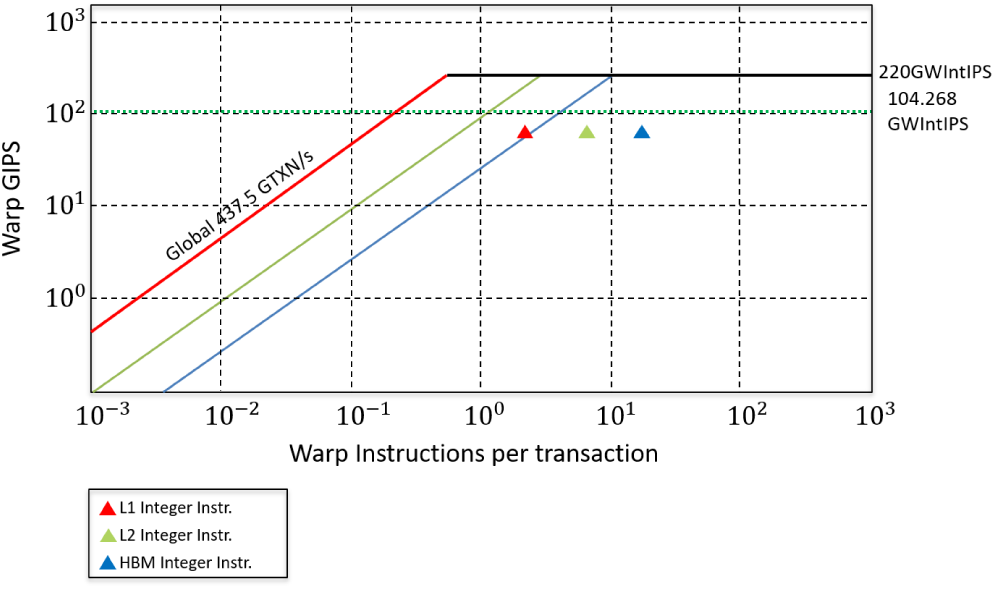
Roofline model plot for a kernel with 64 threads per block and 120000 parallel alignments

**Fig. 5.**
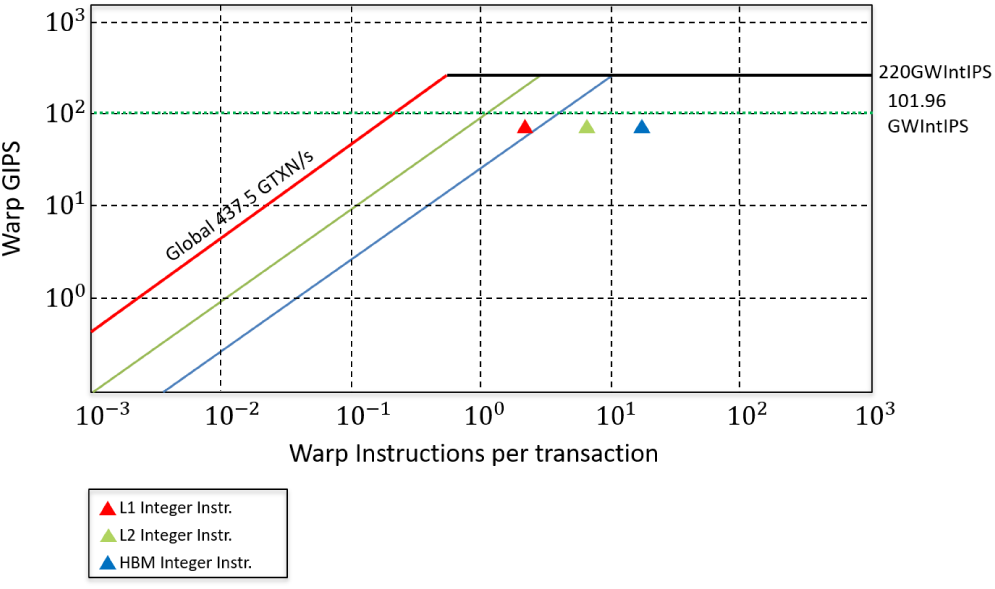
Roofline model plot for a kernel with 128 threads per block and 40000 parallel alignments

**Fig. 6.**
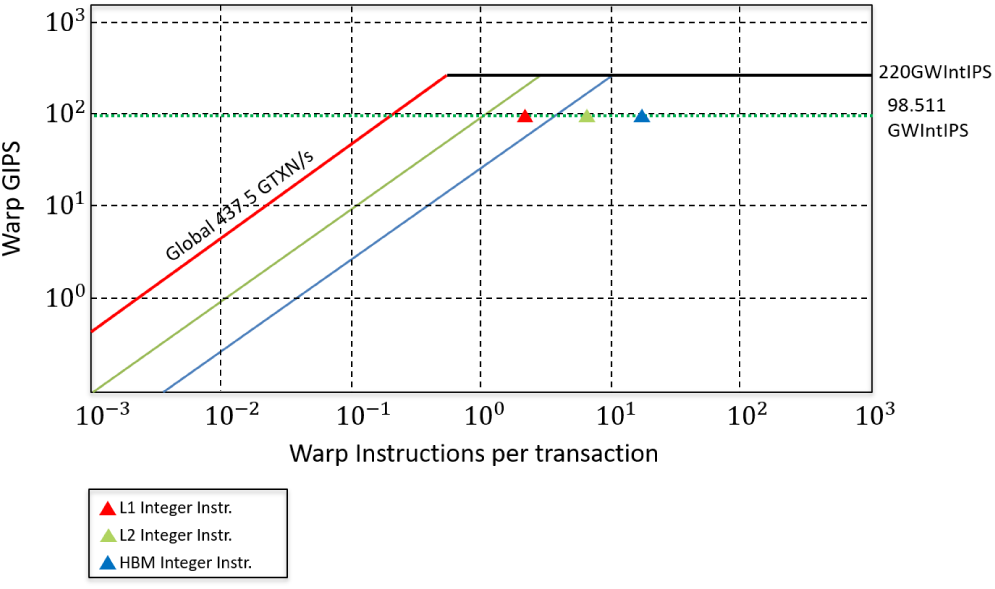
Roofline model plot for a kernel with 255 threads per block and 10900 parallel alignments

## V. CONSENT Integration

The CONSENT tool as presented in [3] contains an error correction module that takes as input a set of raw reads and outputs their corrected version. To do so, the tool first creates a series of overlaps between the reads using minimap2 [19]. At the end of this step, the resulting piles of reads are processed to create a consensus sequence. A segmentation strategy based on the individuation of collinear k-mers is used to subdivide the original MSA task into a series of smaller tasks. This has been shown to have a positive effect on the overall quality of the correction, while reducing the runtime of the correction procedure. One of the most important aspects of this technique is that by transforming the task of a single, very large multiple sequence alignment into a set of smaller MSAs, it changes the requirements of a corresponding GPU acceleration of this procedure: we need a kernel that is optimized to perform a high number of smaller MSA tasks in parallel. For this reason, our GPU implementation is optimized to process small sequences (¡= 255 bp) but a very high number of alignments in parallel (up to 150000). We have empirically verified on real datasets that on average our GPU implementation covers more than 90% of the required alignments for a correction run with default parameters. For the cases where the alignment to perform does not fit on the device, we fall back to running the alignment on CPU. To effectively integrate our GPU kernel within CONSENT, we had to apply some important changes to the correction procedure, as well as create an appropriate software infrastructure to ensure the maximum utilization of our GPU kernel.

### A. Task Batching

In the original CONSENT software, a thread pooling strategy is employed. Each read to correct represents a task that is assigned to an available thread, which performs the whole correction procedure, including the segmentation of the read pile into smaller windows, the corresponding MSA tasks and the computation of the final consensus sequence. To fully exploit the capability of our GPU kernel, we needed to have more alignments to perform in parallel than just the ones corresponding to a single read. To this end, we created a batching strategy that divides the correction procedure in three phases: pre-processing and segmentation, PO graph alignment, and post-processing and consensus computation. For each batch of reads to correct we use multiple threads to perform the first phase for all the reads, then the PO alignment for all the resulting MSA tasks, and lastly we compute the consensus sequence for all the reads. In this way, by selecting an appropriate batch size, we can perform several thousands alignments in parallel.

### B. Kernel Selection

The number of GPU blocks that we can schedule for each kernel call depends on the amount of memory space required by each alignment in the batch. For this reason, our GPU kernels are templates which can be called with different combinations of parameters to ensure the maximum level of parallelism is achieved. We have empirically derived a few optimal combinations for different MSA tasks depending on the number of sequences in a window and the maximum length of the sequences to align. To be able to fully utilize this feature, all the MSA tasks must be divided based on this characteristics and processed by a set of kernels called with the appropriate parameters.

### C. Queue infrastructure

Our GPU-accelerated version of CONSENT relies on a thread-safe queue infrastructure to assign the MSA tasks to the appropriate GPU kernels and manage the pre-processing and post-processing for each read. At the start of the computation, an executor thread is initialized. This thread takes care of calling the GPU MSA procedure whenever a batch of alignment tasks is ready to be processed. When the processing of a batch of tasks is initiated, all the remaining CPU thread are employed with a round robin scheduling policy to perform the pre-processing and segmentation for a pile of reads. After the segmentation, each thread assigns to each of the resulting MSA tasks a label corresponding to an existing kernel template that is the most appropriate to execute PO graph alignment on that specific set of sequences. Immediately after, each thread enqueues the tasks to separate thread-safe queues. Whenever a queue reaches a certain capacity, the tasks are transferred to the GPU and the appropriate set of kernels for PO alignment is called by the executor thread. When all the reads in a CPU batch have been processed, if any alignment tasks remain in any of the queues, they are computed. When all the alignments have been computed, the post processing phase is initiated. All the available CPU threads are again used in a round robin fashion to retrieve the MSA results for each task, compute the consensus sequence for the given task and finally reconstruct the complete corrected read from the fragmented consensus sequences. The process is repeated until all the read piles have been processed. A schematic of this infrastructure is shown in figure 7.

**Fig. 7.**
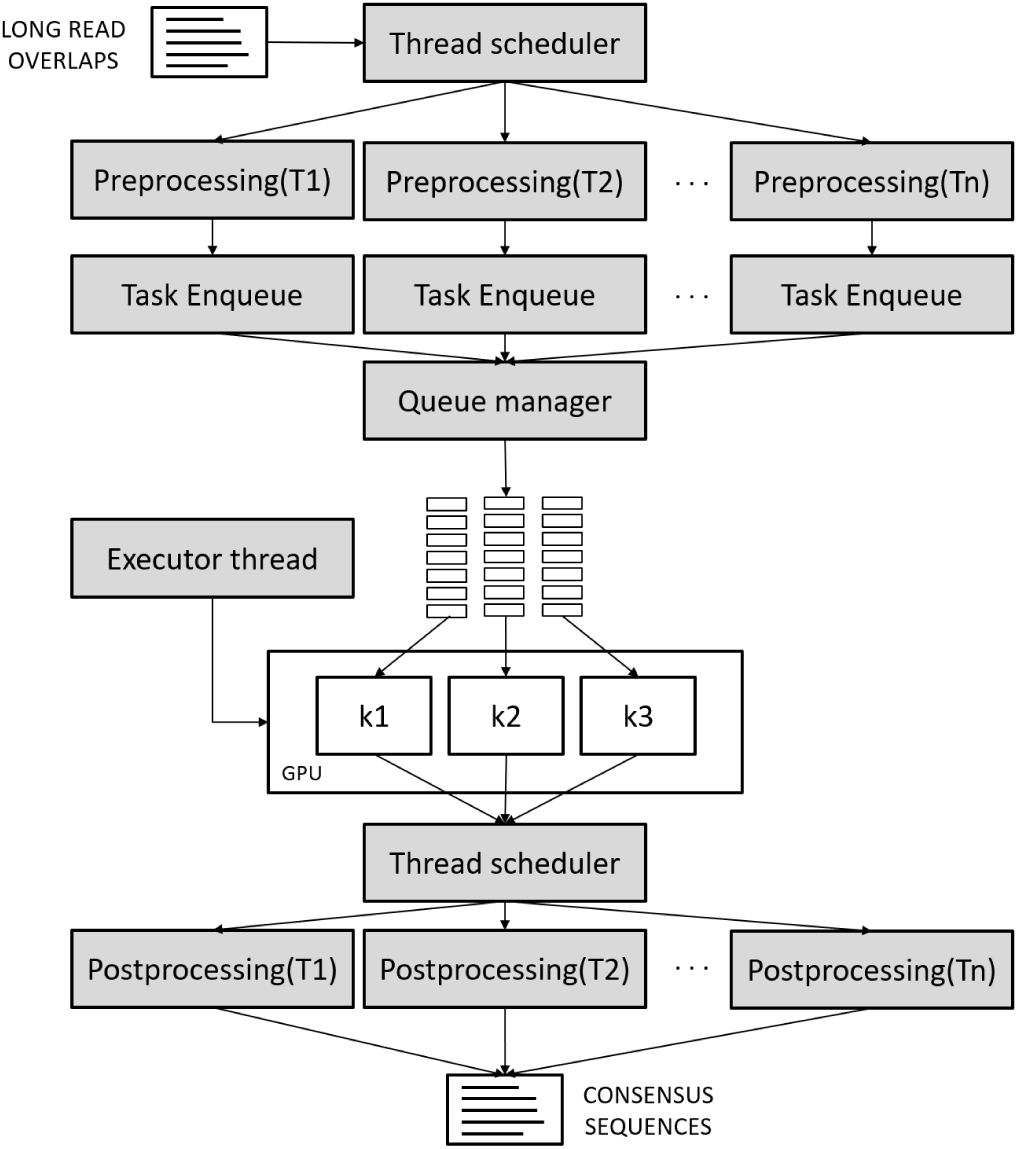
Schematic of the thread-safe queue infrastructure that integrates our GPU kernel in CONSENT

## VI. Discussion

To validate our approach, we compared the performance of our kernel both with the original BOA library and with other implementations of PO alignment on GPU. Moreover, we have evaluated the performance of our integration with CONSENT, by comparing the runtimes of the correction portion of the tool on different datasets.

### A. BOA comparison

Since the target for our implementation is the alignment of small sequences, as a first benchmark we have compared the performance of our GPU implementation to the performance of the BOA software library in a multi-threading environment for a set of randomly generated windows of sequences. The results are shown in table I. In table II we compare our kernel performance to a different CPU architecture against both a single thread and multi-thread CPU implementation. The results show that we can obtain a speedup of up to 3.12x versus 80 threads run on two Intel Xeon Gold 6148 (‘Skylake’) processors running at 2.40 GHz, and up to 6.49x versus 64 threads run on two 2.3 GHz 16-core Intel Xeon Processors E5-2698 v3. These results show that our GPU implementation significantly outperforms state-of-the-art multiprocessor nodes.

**TABLE I.**
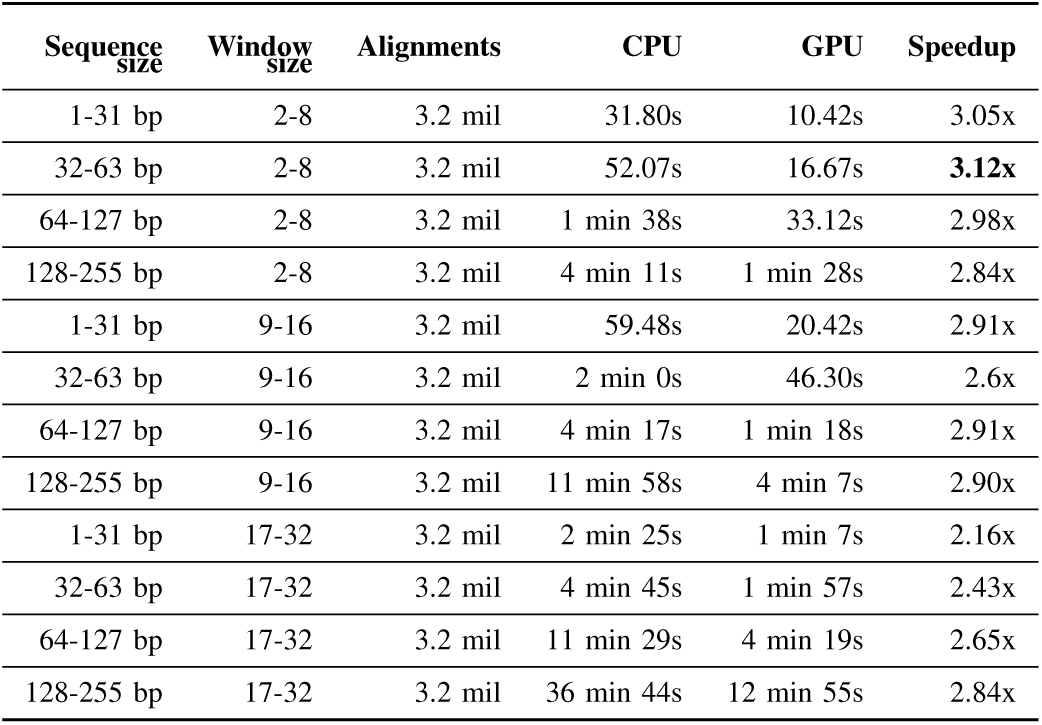
Performance comparison of the PO alignment kernel executed on a NVIDIA Tesla V100 against the CPU implementation of the BOA library executed with 80 parallel threads on two Intel Xeon Gold 6148 (‘Skylake’) running at 2.40 GHz. Both were executed on 3.2 million windows of sequences.

| Sequence size | Window size | Alignments | CPU | GPU | Speedup |
| --- | --- | --- | --- | --- | --- |
| 1-31 bp | 2-8 | 3.2 mil | 31.80s | 10.42s | 3.05x |
| 32-63 bp | 2-8 | 3.2 mil | 52.07s | 16.67s | **3.12x** |
| 64-127 bp | 2-8 | 3.2 mil | 1 min 38s | 33.12s | 2.98x |
| 128-255 bp | 2-8 | 3.2 mil | 4 min 11s | 1 min 28s | 2.84x |
| 1-31 bp | 9-16 | 3.2 mil | 59.48s | 20.42s | 2.91x |
| 32-63 bp | 9-16 | 3.2 mil | 2 min 0s | 46.30s | 2.6x |
| 64-127 bp | 9-16 | 3.2 mil | 4 min 17s | 1 min 18s | 2.91x |
| 128-255 bp | 9-16 | 3.2 mil | 11 min 58s | 4 min 7s | 2.90x |
| 1-31 bp | 17-32 | 3.2 mil | 2 min 25s | 1 min 7s | 2.16x |
| 32-63 bp | 17-32 | 3.2 mil | 4 min 45s | 1 min 57s | 2.43x |
| 64-127 bp | 17-32 | 3.2 mil | 11 min 29s | 4 min 19s | 2.65x |
| 128-255 bp | 17-32 | 3.2 mil | 36 min 44s | 12 min 55s | 2.84x |

**TABLE II.** Performance comparison of the PO alignment kernel executed on a NVIDIA Tesla V100 against the CPU implementation of the BOA library executed on a single thread and with 64 parallel threads on two 2.3 GHz 16-core Intel Xeon Processors E5-2698 V3 with a total of 64 hardware threads. Both were executed on 1.2 million windows of sequences.

| Sequence size | Window size | Alignments | CPU | 1 thread speedup | 64 threads speedup |
| --- | --- | --- | --- | --- | --- |
| 1-31 bp | 2-8 | 1.2 mil | 1 min 34s | 35.31x | 2.6x |
| 32-63 bp | 2-8 | 1.2 mil | 7 min 52s | 82.15x | 3.5x |
| 64-127 bp | 2-8 | 1.2 mil | 26 min 45s | 121.28x | 4.3x |
| 128-255 bp | 2-8 | 1.2 mil | 1 h 42 min | **192.13x** | **6.49x** |

### B. Clara Genomics comparison

To the best of our knowledge, the only other GPU implementation of PO alignment at present is the POA module contined in the Clara Genomics Analysys SDK. To compare our implementation we run both kernels on an NVIDIA Tesla V100 on the same dataset of 2 milion randomly generated windows of sequences ranging from 1 to 255 bp with windows of up to 32 sequences. This choice is motivated by the fact that our kernel targets a specific use case, namely the CONSENT integration. We evaluated on several real datasets for long reads error correction that after the segmentation strategy applied by CONSENT on the initial long reads overlaps over 90% of the resulting windows fall within these dimensions. Therefore we have optimized our kernel to deal efficiently with a very large number of smaller windows of sequences and it is significant to compare to existing implementations through a similar use case. Table III shows the comparison against the Clara Genomics kernel. On the first line the kernel was run with a single cuda stream in MSA generation mode, since our implementation also generates the MSA as output. In this case our implementation outperforms the Clara Genomics PO alignment by 5.89x. In the second line the kernel was run with the Clara Genomics multi-batch benchmark, which uses multiple cuda streams to fill up the GPU. This benchmark outputs the consensus sequence instead of the MSA, since the consensus generation is more optimized than the MSA generation in Clara Genomics. Nonetheless, our kernel achieves a speedup of 3.92x.

**TABLE III.** Performance comparison of the PO alignment kernel executed on a NVIDIA Tesla V100 against the Clara Genomics Analysis SDK implementation of PO alignment. Both were executed on 2 million windows of sequences.

| Sequence size | Window size | Alignments | Our kernel | Clara Genomics | Speedup |
| --- | --- | --- | --- | --- | --- |
| 1-255 bp | 1-32 | 2 mil | 2 min 40s | 15 min 42s | **5.89x** |
| 1-255 bp | 1-32 | 2 mil | 2 min 40s | 10 min 28s | 3.92x |

### C. CONSENT integration evaluation

To evaluate the performance of our integration with CONSENT we have run both the original version presented in [3] and our GPU accelerated version on multiple raw reads datasets. In table IV we report the speedups obtained with our GPU integration. The quality of the results is unchanged, since our kernel performs the same MSA procedure as its CPU counterpart and the corrected reads we obtain from our GPU accelerated version of consent are identical to the ones obtained with the CPU-only version. The performance vary depending on the organism, genome size and coverage, since the number of windows produced as well as the size of the corresponding alignments is heavily dependent on the data. Although the PO alignment is just a portion of the CONSENT error correction procedure, we have been able to obtain significant overall speedup since we are efficiently using both the GPU and CPU resources available on our test system. Moreover, the batching strategy adopted to improve the throughput of the GPU kernel improves the overall performance of CONSENT, at the cost of more memory consumption proportional to the size of the tasks present in a single batch. This result shows how our GPU implementation can be integrated into an error correction tool and improve its overall performance. Given that CONSENT is able to obtain good quality results for a variety of raw read datasets at the cost of a fairly long run time, we have shown how the use of a specific accelerator can improve performance and speedup many genome analysis pipelines that rely on an error correction module to improve the quality of third generation sequencing data.

**TABLE IV.** Performance comparison of CONSENT and our GPU accelerated version. Both software were run on two Intel Xeon Gold 6148 (‘Skylake’) running at 2.40 GHz with 80 parallel threads.

| Dataset | Organism | Dataset size | CONSENT-GPU | CONSENT | Speedup |
| --- | --- | --- | --- | --- | --- |
| SRR10326407 | E. Coli(30x) | 151 Mbp | 6 min 29s | 34 min 36s | 5.3x |
| SRR10326407 | E. Coli(60x) | 290 Mbp | 16 min 44s | 2h 26 min | **8.5x** |
| SRR7743079 | D. Melanogaster(20x) | 2.9 Gbp | 2h 53 min | 6h 17min | 2.18x |
| ERR3454401 | S. Cerevisiae(30x) | 386 Mbp | 1h 6 min | 23 min | 2.86x |
| ERR3454401 | S. Cerevisiae(60x) | 756 Mbp | 3h 0 min | 1h 32 min | 1.95x |

## VII. Conclusions

We presented a GPU accelerated algorithm for multiple sequence alignment based on partial order graphs. The proposed implementation outperforms the state-of-the-art CPU-based POA v2 alignment library for the targeted range of sequence and window lengths, achieving a speedup that ranges from 2.16x to 6.49x versus state-of-the-art CPU nodes run with the optimal number of threads. To evaluate the quality of the proposed GPU implementation, we have devised an extension of the Roofline model for GPU that is able to take into account the nature of the parallelism that characterizes a generic alignment algorithm based on dynamic programming and we show that our kernel achieves near-optimal performance with respect to the performance ceiling computed by our model.

We have also compared our kernel to the PO alignment kernel present in the NVIDIA Clara Genomic Analysis SDK [18] which is, to the best of our knowledge, the only other GPU accelerated multiple sequence alignment based on partial order graphs. We have shown that for the target range of sequences and window lengths we outperform the Clara Genomics PO alignment module by 5.89x on the NVIDIA Tesla V100 GPU. To showcase the practical utility of our alignment kernel we have integrated it into the CONSENT software for long reads self-correction. This use case shows how an alignment strategy based on the segmentation of longer overlaps between long reads into smaller alignment windows presents an opportunity to exploit the parallel processing power offered by GPU architectures. We have created the appropriate software infrastructure to ensure that we fully take advantage of the computational capabilities of the GPU while also exploiting the CPU resources as much as possible. We have shown that we maintain the quality of the results while reducing the execution time of the error correction phase by two to four times for real raw read datasets for different organisms.

## Acknowledgements

We would like to thank Alberto Zeni, Guido Walter Di Donato and Muaaz Awan for useful suggestions and valuable discussions. This work is supported by the Advanced Scientific Computing Research (ASCR) program within the Office of Science of the DOE under contract number DE-AC02-05CH11231. This research was also supported by the Exascale Computing Project (17-SC-20-SC), a collaborative effort of the U.S. Department of Energy Office of Science and the National Nuclear Security Administration. We used resources of the NERSC supported by the Office of Science of the DOE under Contract No. DEAC02-05CH11231.

## References

[1] H. Zhang, C. Jain, and S. Aluru, “A comprehensive evaluation of long read error correction methods,” BioRxiv, p. 519330, 2019.

[2] R. Vaser, I. Sović, N. Nagarajan, and M. Šikić, “Fast and accurate de novo genome assembly from long uncorrected reads,” Genome research, vol. 27, no. 5, pp. 737–746, 2017.

[3] P. Morisse, C. Marchet, A. Limasset, T. Lecroq, and A. Lefebvre, “Consent: Scalable self-correction of long reads with multiple sequence alignment,” BioRxiv, p. 546630, 2019.

[4] A. Limasset, “Boa repository,” 2018, https://github.com/Malfoy/BOA [Accessed: 2019-10-28].

[5] C. Lee, C. Grasso, and M. F. Sharlow, “Multiple sequence alignment using partial order graphs,” Bioinformatics, vol. 18, no. 3, pp. 452–464, 2002.

[6] N. Ding and S. Williams, “An instruction roofline model for gpus,” 2019 IEEE/ACM Performance Modeling, Benchmarking and Simulation of High Performance Computer Systems (PMBS), 2019.

[7] S. F. Altschul, T. L. Madden, A. A. Schäffer, J. Zhang, Z. Zhang, W. Miller, and D. J. Lipman, “Gapped blast and psi-blast: a new generation of protein database search programs,” Nucleic acids research, vol. 25, no. 17, pp. 3389–3402, 1997.

[8] S. R. Eddy, “Profile hidden markov models.” Bioinformatics (Oxford, England), vol. 14, no. 9, pp. 755–763, 1998.

[9] L. Wang and T. Jiang, “On the complexity of multiple sequence alignment,” Journal of computational biology, vol. 1, no. 4, pp. 337–348, 1994.

[10] J. D. Thompson, D. G. Higgins, and T. J. Gibson, “Clustal w: improving the sensitivity of progressive multiple sequence alignment through sequence weighting, position-specific gap penalties and weight matrix choice,” Nucleic acids research, vol. 22, no. 22, pp. 4673–4680, 1994.

[11] C.-L. Hung, Y.-S. Lin, C.-Y. Lin, Y.-C. Chung, and Y.-F. Chung, “Cuda clustalw: An efficient parallel algorithm for progressive multiple sequence alignment on multi-gpus,” Computational biology and chemistry, vol. 58, pp. 62–68, 2015.

[12] Y. Liu, B. Schmidt, and D. L. Maskell, “Msa-cuda: multiple sequence alignment on graphics processing units with cuda,” in 2009 20th IEEE International Conference on Application-specific Systems, Architectures and Processors. IEEE, 2009, pp. 121–128.

[13] C. Notredame, D. G. Higgins, and J. Heringa, “T-coffee: A novel method for fast and accurate multiple sequence alignment,” Journal of molecular biology, vol. 302, no. 1, pp. 205–217, 2000.

[14] J. Blazewicz, W. Frohmberg, M. Kierzynka, and P. Wojciechowski, “G-msa—a gpu-based, fast and accurate algorithm for multiple sequence alignment,” Journal of Parallel and Distributed Computing, vol. 73, no. 1, pp. 32–41, 2013.

[15] C. Lee, “Generating consensus sequences from partial order multiple sequence alignment graphs,” Bioinformatics, vol. 19, no. 8, pp. 999–1008, 2003.

[16] M. Blanchette, W. J. Kent, C. Riemer, L. Elnitski, A. F. Smit, K. M. Roskin, R. Baertsch, K. Rosenbloom, H. Clawson, E. D. Green et al., “Aligning multiple genomic sequences with the threaded blockset aligner,” Genome research, vol. 14, no. 4, pp. 708–715, 2004.

[17] B. Paten, D. Earl, N. Nguyen, M. Diekhans, D. Zerbino, and D. Haussler, “Cactus: Algorithms for genome multiple sequence alignment,” Genome research, vol. 21, no. 9, pp. 1512–1528, 2011.

[18] NVIDIA, “Clara genomics sdk,” 2019, https://developer.nvidia.com/Clara-Genomics [Accessed: 2019-10-28].

[19] H. Li, “Minimap2: pairwise alignment for nucleotide sequences,” Bioinformatics, vol. 34, no. 18, pp. 3094–3100, 2018.

